# The CIC-ERF co-deletion underlies fusion independent activation of ETS family member, ETV1, to drive prostate cancer progression

**DOI:** 10.1101/2022.01.26.477820

**Authors:** Nehal Gupta, Hanbing Song, Wei Wu, Rovingaile Kriska Ponce, Yone Kawe Lin, Ji Won Kim, Eric J Small, Felix Y. Feng, Franklin W. Huang, Ross A. Okimoto

## Abstract

The dysregulation of ETS family transcription factors drives human prostate cancer. The majority of prostate cancer is the result of chromosomal rearrangements that lead to aberrant ETS gene expression. The mechanisms that lead to fusion independent ETS factor upregulation and prostate oncogenesis remain unknown. Here, we show that two neighboring transcription factors, Capicua (*CIC*) and ETS2 repressor factor (*ERF*), which are co-deleted in human prostate tumors can drive prostate oncogenesis. Concurrent *CIC* and *ERF* loss commonly occurs through focal genomic deletions at chromosome 19q13.2. Mechanistically, *CIC* and *ERF* co-bind the proximal regulatory element and mutually repress the ETS transcription factor, *ETV1*. Targeting ETV1 in *CIC* and *ERF* deficient prostate cancer limits tumor growth. Thus, we have uncovered a fusion independent mode of ETS transcriptional activation defined by concurrent loss of *CIC* and *ERF*.

## Introduction

Prostate cancer (PCa) is the most common solid tumor malignancy in men. Activation of ETS transcription factors, *ERG, ETV1, ETV4*, and *ETV5*, are present in approximately 60% of PCa, underscoring their importance in prostate oncogenesis ^1^. In human PCa, ETS transcription factors are most commonly activated through gene rearrangements that fuse the androgen responsive gene, *TMPRSS2*, to either *ERG, ETV1, ETV4*, or *ETV5* ^2^. Beyond ETS transcription factor fusions, little is known about the underlying molecular mechanisms that lead to increased expression of wild-type ETS factors, which confer aggressive malignant phenotypes and associate with poor clinical outcomes in fusion negative PCa patients ^3^.

Capicua (CIC) is an HMG box transcription factor that silences *ETV1, ETV4*, and *ETV5* through direct target gene repression ^4,5^. CIC is frequently altered in human cancer, where it functionally suppresses tumor growth and metastasis ^5–13^. Notably, in PCa, *CIC* is commonly altered through genomic loss (homozygous and heterozygous deletion) in ∼10% of PCa patients ^14–19^ and inactivation of *CIC* de-represses *ETV1, ETV4*, and *ETV5* transcription to promote tumor progression. Leveraging mutational profiling data from multiple PCa cohorts, we previously observed concurrent loss of the ETS2 repressor factor (ERF) in CIC-deficient prostate tumors ^19^. Combinatorial loss is most commonly the result of focal deletions (homozygous and heterozygous) at the 19q13.2 locus, where *CIC* and *ERF* are physically adjacent (long and short isoforms of *CIC* are separated from *ERF* by approximately 15 and 30kb, respectively) to one another in the genome. Since ERF is a transcriptional repressor that binds ETS DNA motifs ^20^, we hypothesized that in a fusion independent manner, CIC and ERF cooperate to mutually suppress ETS target genes in prostate cancer.

Through an integrative genomic and functional analysis, we mechanistically show that CIC and ERF directly bind and co-repress a proximal *ETV1* regulatory element limiting prostate cancer progression. Concomitant loss of CIC and ERF de-represses *ETV1* mediated transcriptional programs and confer *ETV1* dependence in multiple *in vitro* and *in vivo* prostate cancer models. Thus, we reveal a fusion independent mechanism to derepress ETS-mediated prostate cancer progression and uncover a therapeutic approach to target CIC-ERF co-deleted prostate cancer.

## Results

CIC is a transcription factor (TF) that directly suppresses *PEA3* (*ETV1, ETV4*, and *ETV5*) TF family members ^5,6,21,22^. CIC silences target genes through direct binding of a highly conserved DNA binding motif ((T(G/C)AAT(G/A)AA) (Fig 1A) ^21,23^. CIC is commonly altered in multiple human cancer subtypes where it suppresses tumor growth and metastasis ^5^. *CIC* is located on chromosome 19q13.2, directly adjacent to another transcriptional repressor, namely the ETS2 repressor factor (*ERF*) (Fig 1b). ERF binds and competes for ETS TF binding sites (GGAA-motifs) and is frequently altered in human prostate cancer, predominantly through focal deletions ^20,24^. We thus hypothesized that concurrent loss of *CIC* and *ERF* may de-repress an ETS driven transcriptional program that drives PCa progression in a fusion independent manner. To explore this, we first queried 15 independent prostate cancer datasets curated on cBioPortal ^25,26^. We analyzed over 6047 prostate cancer tumors from 5839 patients and identified a high co-occurrence rate *(p* < 0.001, two-sided Fisher’s exact test (FET)) for *CIC* (10%) and *ERF* (12%) homozygous and heterozygous deletions (Fig. 1c), suggesting that concurrent loss occurs through focal copy number change at the 19q13.2 locus. Through analysis of these clinically annotated specimens, we observed that the CIC-ERF co-deletion was present at an increased frequency in prostate cancers with higher Gleason Scores and later tumor stages when compared to *CIC-ERF* replete tumors (Fig 1d). In order to understand the association between *CIC* and/or *ERF* alterations in specific prostate cancer cohorts, we stratified published datasets to identify patients that represent primary prostate cancer ^14,16,27^ and metastatic castrate resistant prostate cancer (mCRPC) ^15,17,18^. This analysis revealed enrichment of *CIC* and *ERF* alterations including the *CIC-ERF* co-deletion in mCRPC samples (Fig. 1e). Importantly, *CIC-ERF* co-deleted tumors clustered as a distinct subgroup when compared to the more well-characterized molecular subsets including *ERG, ETV1, ETV4, SPOP*, and *FOXA1* altered prostate cancers suggesting a new subtype of prostate cancer (Fig 1f). In order to explore clinical outcomes of patients harboring *CIC-ERF* co-deleted tumors, we performed a survival analysis using the aforementioned prostate cancer datasets and observed significantly worse outcomes in patients that harbored the *CIC-ERF* co-deletion (*p* = 0.001, Disease-free survival and p = 0.01, Progression-free survival) (Fig 1g). Collectively, these findings indicated that *CIC* and *ERF* are co-deleted with increasing frequency in mCRPC and that *CIC-ERF* co-deletion is associated with worse clinical outcomes in prostate cancer patients.

**Figure 1.**
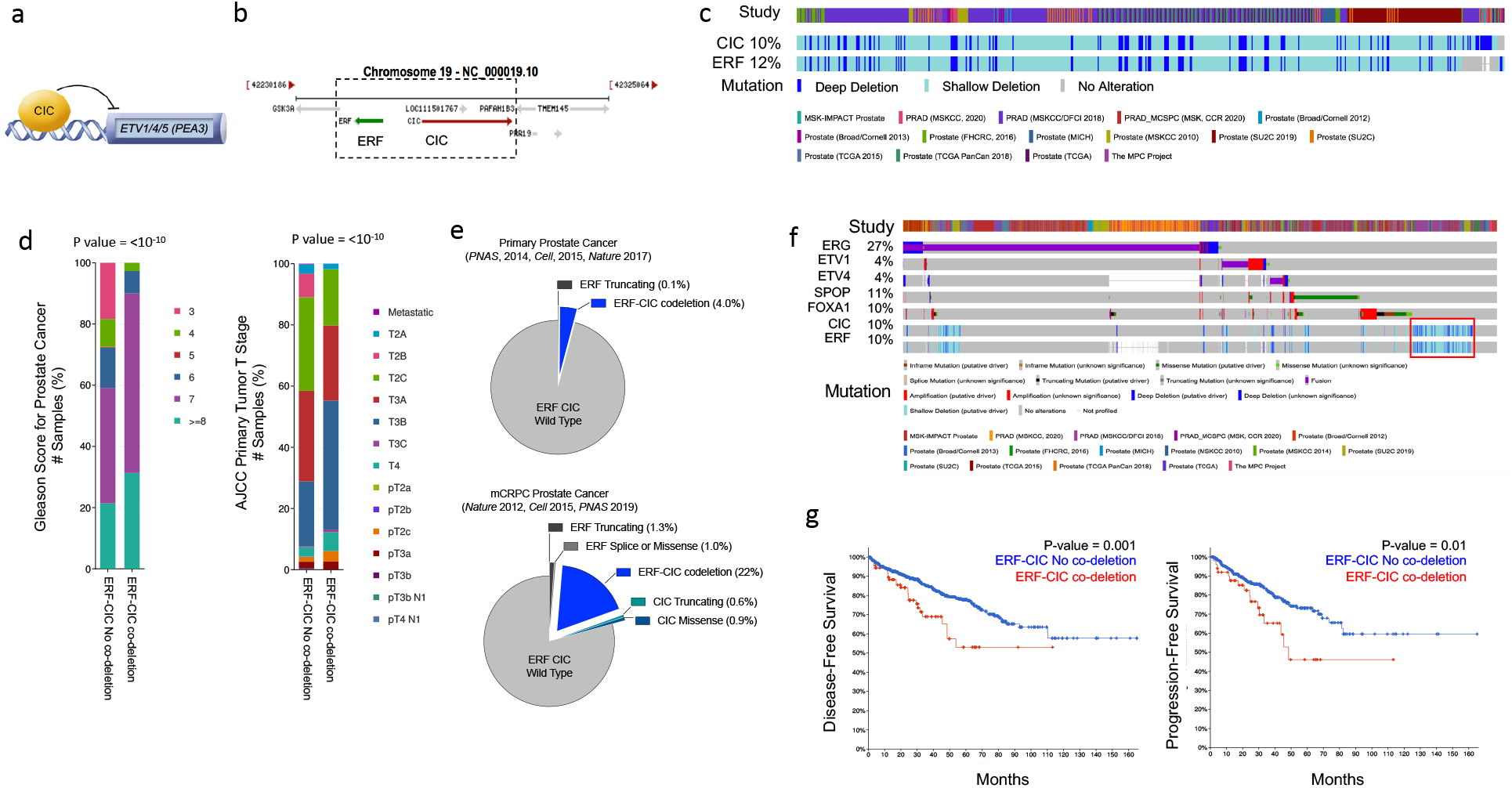
CIC and ERF are co-deleted in aggressive prostate cancer and associate with worse clinical outcomes. (a) CIC transcriptionally represses ETV1/4/5. (b) The 19q13.2 genomic locus demonstrating the physical location of ERF and CIC. (c) 15 PCa studies (cBioPortal) demonstrating the co-occurrence of ERF and CIC homozygous and heterozygous deletions. The co-occurrence of ERF and CIC alterations is highly significant (P<0.001 co-occurrence, Fisher exact test). (d) ERF-CIC co-deleted PCa stratified by Gleason Score and Tumor Stage. (e) Frequency of ERF and CIC alterations in primary PCa (top) and mCRPC (bottom), demonstrating enrichment in mCRPC. (f) Onco-print of known genetic drivers (ERG, ETV1, ETV4, SPOP, FOXA1) of PCa aligned with CIC and ERF (cBioPortal). CIC-ERF co-deleted prostate tumors (red box) do not frequently co-occur with other known oncogenic events. (g) Survival analysis performed using 18 prostate cancer datasets from TCGA. DFS and PFS in patients harboring the ERF-CIC codeletion (red) vs. no ERF-CIC co-deletion (blue). P=value, log-rank.

Our initial findings provided rationale to explore the genetic and functional relationship between CIC and ERF. Independently, CIC and ERF have previously been reported to promote malignant phenotypes, including tumor growth and metastasis in multiple human cancer subsets ^6,7,32,33,8,10,19,20,28–31^. Since our clinical data indicated that combinatorial loss of CIC and ERF was associated with worse patient outcomes, we hypothesized that CIC and ERF loss may cooperate to enhance prostate cancer progression. To investigate this, we engineered ERF, CIC, or both CIC and ERF-deficient immortalized prostate epithelial cells (PNT2) and performed a series of *in vitro* and *in vivo* experiments to test the combinatorial effect of the *CIC-ERF* co-deletion. Compared to single gene loss of CIC or ERF, genetic inactivation of both *CIC* and *ERF* increased colony formation (Fig 2a-b, Supplementary Fig 1a) and spheroid formation (Fig 2c-d) in PNT2 cells. Additionally, genetic silencing of both *CIC* and *ERF* increased the frequency of subcutaneous tumor xenograft formation in immunodeficient (SCID) mice compared to control (Fig. 2e, Supplementary Fig 1b). *CIC-ERF* co-deletion also enhanced malignant phenotypes including cellular viability, invasiveness, and migratory capacity in PNT2 prostate epithelial cells (Fig 2f-h). Thus, our findings demonstrate that the combination of CIC and ERF loss augments the transformation of prostate epithelial cells and promotes malignant phenotypes.

**Figure 2.**
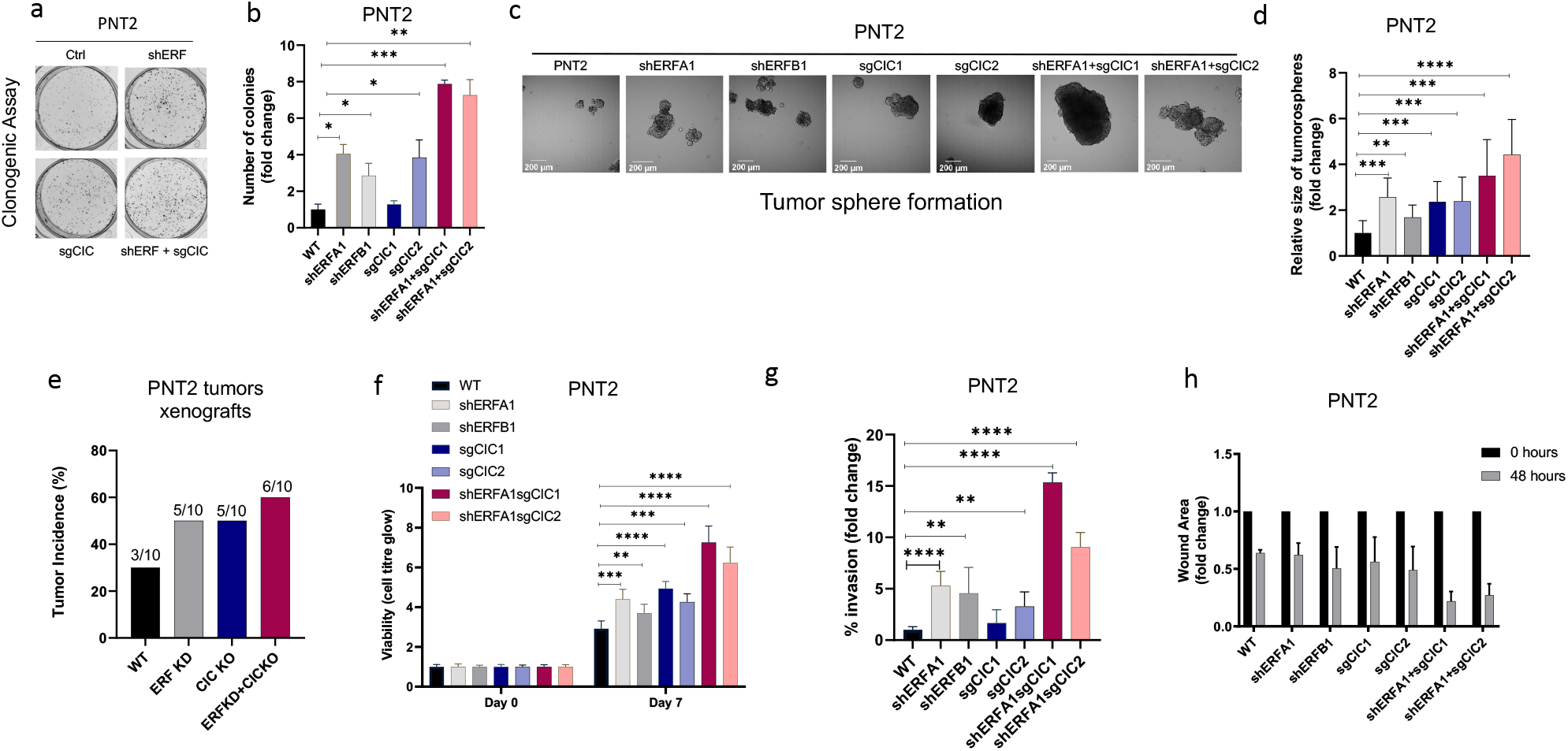
CIC and ERF loss promote tumor formation and control malignant potential in prostate epithelial cells. (a) Clonogenic assay comparing PNT2 cells expressing ERF KD, CIC KO, or ERF KD+CIC KO compared to control. (b) Number of colonies for each condition in (a). (c) Spheroid growth assay using PNT2 cells expressing ERF KD, CIC KO, ERF KD+CIC KO, vs. control. (d) Size of the sphere for each condition in (c). Error bars represent SD. P values were calculated using Student’s t test. *p<0.05, **p<0.01, ***p<0.001, ****p<0.0001. (e) Bar graph comparing the incidence of PNT2 parental (N=3/10), PNT2 ERF KD (5/10), PNT2 CIC KO (N=5/10) or PNT2 ERF KD+CIC KO (N=6/10) tumor formation in immunodeficient mice. (f) Cell-titer glo viability assay, (g) Transwell assay, and (h) wound healing assay comparing PNT2 ERF KD, CIC KO, and ERF KD+CIC KO to control. Error bars represent SD. P values were calculated using Student’s t test. **p<0.01, ***p<0.001, ****p<0.0001.

To assess the functional role of CIC and ERF in the context of human prostate cancer progression, we leveraged two genetically annotated, androgen-insensitive prostate cancer cell lines, DU-145 and PC-3. DU-145 cells harbor a loss-of-function ERF mutation (*ERF*^*A132S*^) ^19^ and express functional wild-type *CIC (*Supplementary Fig 2a). By comparison, PC-3 cells are deficient in CIC (CIC deletion) and retain functional ERF (Supplementary Fig 2b) ^9,25^. Thus, these cell line models provided isogenic systems to functionally interrogate the role of CIC and ERF in human prostate cancer. Specifically, genetic reconstitution of *ERF* into *ERF* deficient DU-145 cells decreased colony formation in both *CIC* proficient (parental cells) and *CIC* knockout (KO) conditions (Fig 3a-b, Supplementary Fig. 2c-d). While *CIC* loss did not enhance colony formation in *ERF* deficient DU-145 cells, it significantly increased tumor cell viability, invasion, and migratory capacity compared to control (Fig 3c-d, Supplementary Fig 2e). Importantly, we observed that ERF expression in *CIC* KO DU-145 cells rescued the CIC-mediated effects on viability and migration/invasion (Fig 3c-d, Supplementary Fig 2e). These findings indicate that rescuing of ERF can partially restore the functional effects of CIC loss in DU-145 cells. To further understand if ERF could suppress tumor growth *in vivo*, we reconstituted wildtype (WT) ERF into CIC proficient and deficient (CIC KO) DU-145 cells and generated subcutaneous xenografts in immunodeficient mice (NU/J). Consistent with our *in vitro* data, CIC KO increased the tumor growth rate *in vivo* and genetic reconstitution of WT ERF partially suppressed tumor growth compared to DU-145 CIC KO cells (Fig 3e). Moreover, genetic reconstitution of ERF into CIC proficient DU-145 cells suppressed the tumor growth rate *in vivo* (Fig 3e). We next used PC-3 cells, a *CIC* deficient prostate cancer cell line that has a homozygous deletion of CIC and expresses functional WT ERF^9,25,26^ to further test how ERF and CIC functionally interact in the context of human prostate cancer. We first noted that ERF overexpression or reconstitution of CIC alone decreased PC-3 colony formation, with combinatorial ERF overexpression and CIC rescue having the most significant reduction compared to parental PC-3 cells (Fig 3f-g, Supplementary Fig 2f-g). Moreover, ERF and CIC expression had a similar impact on decreasing PC-3 viability, invasion and migration capacity (Fig 3h-i, Supplementary Fig 2h). Similarly, genetic loss of *ERF* resulted in significant increase of colony formation, viability and invasion compared to parental PC-3 cells (Fig 3f-i, Supplementary Fig 2i). Interestingly, overexpression of ERF in mice bearing CIC deficient PC-3 tumor xenografts significantly decreased tumor growth compared to PC-3 parental and PC-3 cells expressing WT CIC (genetic rescue of CIC) (Fig 3j). Since we consistently observed that ERF expression could partially rescue the effects of CIC loss in prostate cancer, we hypothesized that WT CIC and ERF potentially cooperate to limit prostate cancer progression.

**Figure 3.**
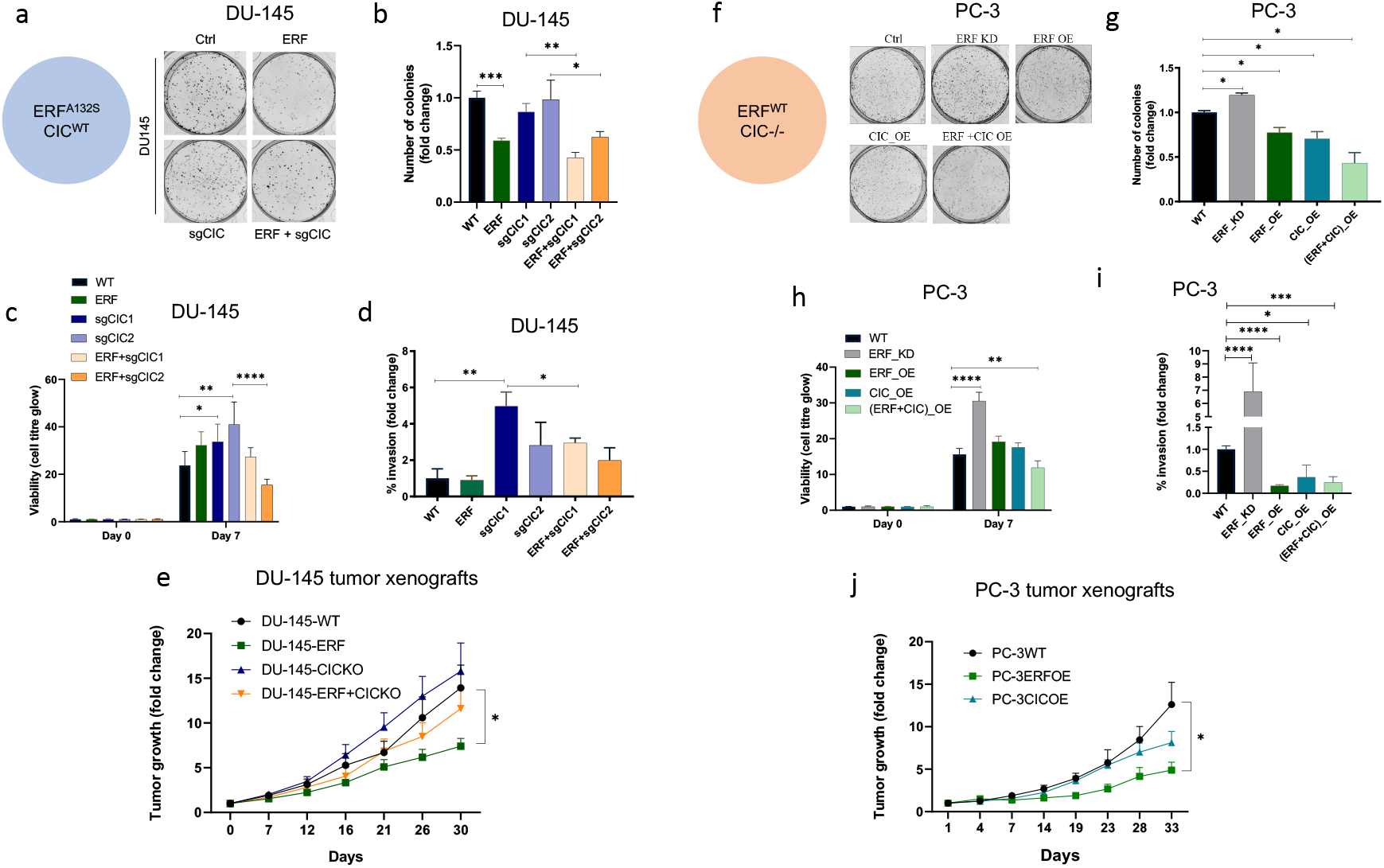
CIC and ERF mutually suppress malignant phenotypes in human prostate cancer. (a) Clonogenic assay in DU-145 cells with ERF rescue, CIC KO, or ERF rescue + CIC KO compared to parental. (b) Number of colonies for each condition in (a). (c) Cell-titer glo viability assay and (d) Transwell assay comparing DU-145 parental cells to DU-145 with ERF rescue, CIC KO, or ERF rescue + CIC KO. P values were calculated using Student’s t test. *p<0.05, **p<0.01, ***p<0.001, ****p<0.0001. Error bars represent SD. (e) Relative tumor volume in mice bearing DU-145 parental, DU-145 ERF, DU-145 with CIC KO, or DU-145 ERF +CIC KO xenografts (N=10). P values were calculated using Student’s t test. *p<0.05. Error bars represent SEM. (f) Clonogenic assay in PC-3 cells expressing ERF KD, ERF OE, CIC OE, or (ERF+CIC) OE compared to control. (g) Number of colonies for each condition in (f). (h) Cell-titer glo viability assay and (i) Transwell assay comparing different groups in PC-3 cells (WT, ERF KD, ERF OE, CIC OE, or (ERF+CIC) OE). P values were calculated using Student’s t test. *p<0.05, **p<0.01, ***p<0.001, ****p<0.0001. Error bars represent SD. (j) Relative tumor volume in mice bearing PC-3 parental cells, PC-3 ERF OE and PC-3 CIC OE over time (N=10). P values were calculated using Student’s t test. *p<0.05. Error bars indicate SEM.

In order to mechanistically define how CIC and ERF (two transcription factors with known repressor function) were interacting to functionally regulate prostate cancer, we performed chromatin immunoprecipitation followed by sequencing (ChIP-Seq) using a validated CIC antibody ^6,28,34^ in PNT2 prostate epithelial cells and compared this to a publicly available ERF ChIP-Seq dataset in VCaP prostate cancer cells ^20^ (we were unsuccessful at pulling down ERF in PNT2 cells). This analysis identified 178 high confidence (FDR <= 0.05) CIC peaks that mapped to 130 annotated genes including known targets, *ETV1, ETV4*, and *ETV5*. Globally, CIC peaks were localized to distinct genomic regions including: promoters (38.6%), UTRs (1.14%), introns (19.9%), distal intergenic regions (38.64%). Interestingly, the distribution of ERF peaks were similar to CIC, with 32.8% in promoters, 1.2% in UTRs, 29.2% intronic, and 33.8% in distal intergenic regions (Fig 4a). Next, through a comparative ChIP-Seq analysis (Fig 4b), we identified 91 shared CIC and ERF target genes. Importantly, we focused on genes with shared CIC and ERF binding sites to potentially explain the functional cooperativity that we observed in our prior studies. In order to narrow down potential candidates, we performed Functional Clustering Analyses using the 91 shared CIC and ERF target genes (Supplementary Table 1). Among these putative CIC and ERF targets, the *PEA3* (*ETV1, ETV4*, and *ETV5*) TFs (known oncogenic drivers in prostate cancer) ^3,35,36^ were found to be the most highly enriched family. These findings suggested that CIC and ERF may co-regulate PEA3 family members through direct transcriptional control. In order to further identify how CIC and/or ERF impact *PEA3* TF expression, we performed quantitative RT-PCR (qRT-PCR), assessing *ETV1, ETV4*, or *ETV5* mRNA levels in response to genetic silencing of CIC and/or ERF. As expected, *CIC* KO and/or combinatorial *CIC* and *ERF* loss in PNT2 cells (ERF and CIC WT) consistently increased *ETV1, ETV4, and ETV5* levels compared to control (Supplementary Fig 3a-f). In contrast, genetic silencing of *ERF* consistently increased *ETV1* mRNA expression, but not *ETV4* or *ETV5* (Fig. 4c, Supplementary Fig 3g-j). These findings indicated that *ETV1* may be a shared transcriptional target of CIC and ERF in prostate cells. To confirm CIC and ERF binding to *ETV1*, we first localized putative CIC (TGAATGGA) and ERF (GGAA) DNA binding sites within the proximal upstream regulatory element of *ETV1* and independently confirmed CIC and ERF occupancy of the *ETV1* promoter through ChIP-PCR (Fig 4d-g). To extend these findings into the context of prostate cancer, we reconstituted *ERF* in *ERF* deficient DU-145 cells and this consistently decreased *ETV1* expression (not *ETV4* or *ETV5*) in both CIC proficient and CIC deficient settings (Fig 4h, Supplementary Fig 3k-n). Moreover, *ERF* KD or *ERF* overexpression in CIC deficient PC-3 cells increased and decreased *ETV1* mRNA expression, respectively (Fig 4i-j). Since *ETV1* is a known target of CIC ^12,21^, we focused on further validating *ETV1* as a molecular target of ERF. To this end, we engineered *ETV1* luciferase based promoter assays and observed a decrease in luciferase activity following ERF expression in 293T and DU-145 cells (Fig 4k-l). These genetic tools further validate that ERF can suppress *ETV1* expression through direct transcriptional silencing of the *ETV1* promoter and identifies ETV1 as a novel target of ERF. Consistent with a repressor function, ERF loss was previously shown to transcriptionally associate with ETV1-regulated gene set signatures ^19^. Yet it remains unclear if ERF can directly regulate *ETV1* and how combinatorial loss of CIC and ERF controls ETV1-mediated (or other ETS family members) transcriptional programs. To explore this, we leveraged our dual CIC and ERF deficient PNT2 cells and performed single-sample Gene Expression Analysis (ssGSEA) ^37^ within The Cancer Genome Atlas prostate cancer (TCGA-PRAD) dataset. We found that CIC and ERF loss were significantly associated with the *ETV1*-regulated gene set (Information coefficient (IC) = 0.619, *p* = 0.0009), but was anti-correlated with the *TMPRSS2-ERG* fusion signature gene set ^38^ (Fig. 4m, IC = -0.45, *p* = 0.0009). Thus, the enrichment of the *ETV1*-regulated gene set signature was shared between *ERF* loss alone ^19^ and *CIC-ERF* dual suppression (Fig 4m). In contrast, combinatorial CIC and ERF loss negatively correlated with the *TMPRSS2-ERG* fusion signature gene set, which was not consistent with our prior studies using ERF KD alone ^19^. These findings led us to hypothesize that the dual suppression of CIC and ERF may increase ETV1-mediated transcriptional programs in prostate cancer. This was supported by two major lines of experimental and conceptual evidence including: 1) dual suppression of *ETV1* expression and ETV1 mediated transcriptional output by CIC and ERF; and 2) the majority of tumors derived from prostate cancer patients that harbor *ERF* deletions also contain deletions in *CIC*.

**Figure 4.**
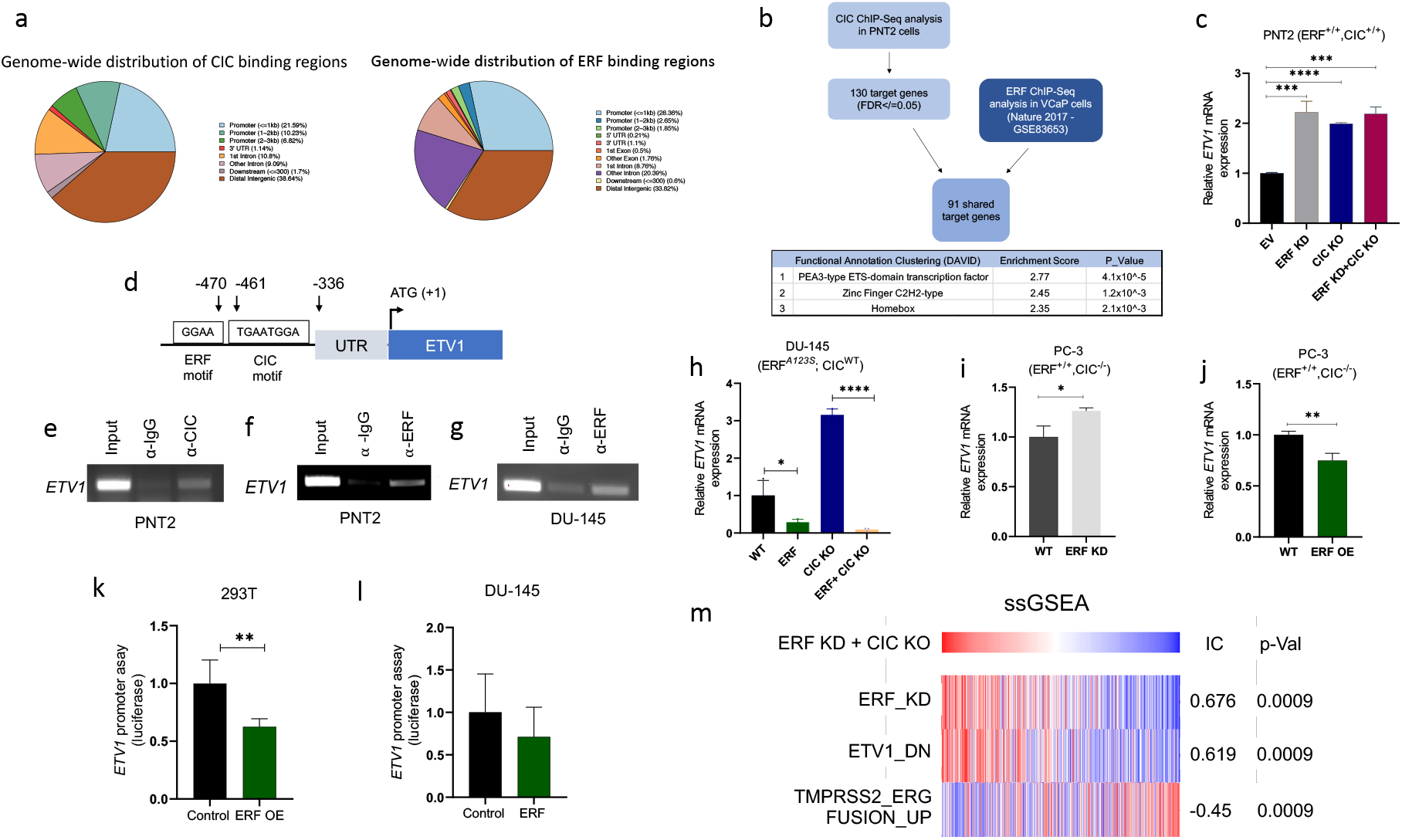
CIC and ERF cooperatively bind an *ETV1* regulatory element to suppress *ETV1* expression and transcriptional activity. (a) Percentage of CIC and ERF peaks located in defined genomic regions. (b) Schematic algorithm to identify shared CIC and ERF target genes in prostate cells (top). Functional Clustering Analysis of the 91 shared CIC and ERF target genes using DAVID (bottom table). (c) *ETV1* mRNA expression in PNT2 (CIC-ERF-replete) cells with ERF KD, CIC KO, or ERF KD + CIC KO. (d) Schematic of CIC and ERF DNA-binding motifs in the *ETV1* promoter. (e) ChIP-PCR from PNT2 cells showing CIC occupancy on the *ETV1* promoter. (f-g) ChIP-PCR with ERF occupancy on the *ETV1* promoter. (h) *ETV1* mRNA expression in DU-145 (ERF-deficient) cells with ERF rescue, CIC KO, or ERF rescue + CIC KO. *ETV1* mRNA expression in PC-3 cells with ERF KD (i) and ERF OE (j). P values were calculated using Student’s t test. *p<0.05, **p<0.01, ****p<0.0001. Error bars represent SD. Performed in triplicate. *ETV1* luciferase promoter assay in 293T (k) and DU-145 (l) cells comparing EV with ERF OE (n = 6). Student’s t test, *p<0.05. Error bars represent SD. (m) Single Sample Gene Set Enrichment Analysis (ssGSEA) alignments comparing gene expression patterns in PNT2 cells with ERF KD and CIC KO. IC = information coefficient.

In order to demonstrate enhanced survival dependence on ETV1 in prostate cancer cells, we genetically silenced *CIC* in ERF-deficient DU-145 cells and assessed drug sensitivity to an ETV1 inhibitor (BRD32048) (Supplementary Fig 4a-c) ^39^. Since BRD32048 was previously shown to decrease invasiveness in ETV1-fusion positive prostate cancer cells (LNCaP), but not significantly impact tumor cell viability, we unexpectedly observed that silencing CIC in DU-145 cells could enhance sensitivity to BRD32048 (Fig 5a). To further confirm these findings and to mitigate potential off-target effects, we silenced *ETV1* using siRNA in DU-145 cells expressing Crispr-based sgRNA targeting CIC (Supplementary Fig 4d-f). Consistent with our chemical inhibitor studies, the viability of DU-145 *CIC* KO cells was decreased upon *ETV1* inhibition (Fig 5b). Similarly, pharmacologic and genetic ETV1 inhibition decreased invasiveness of CIC deficient DU-145 cells (Fig. 5c-d). These findings indicate that loss of CIC in ERF deficient prostate cancer cells can enhance sensitivity to ETV1-directed therapies. Thus, CIC and ERF cooperate to silence ETV1 transcriptional programs, limiting ETV1-mediated prostate cancer progression.

**Figure 5.**
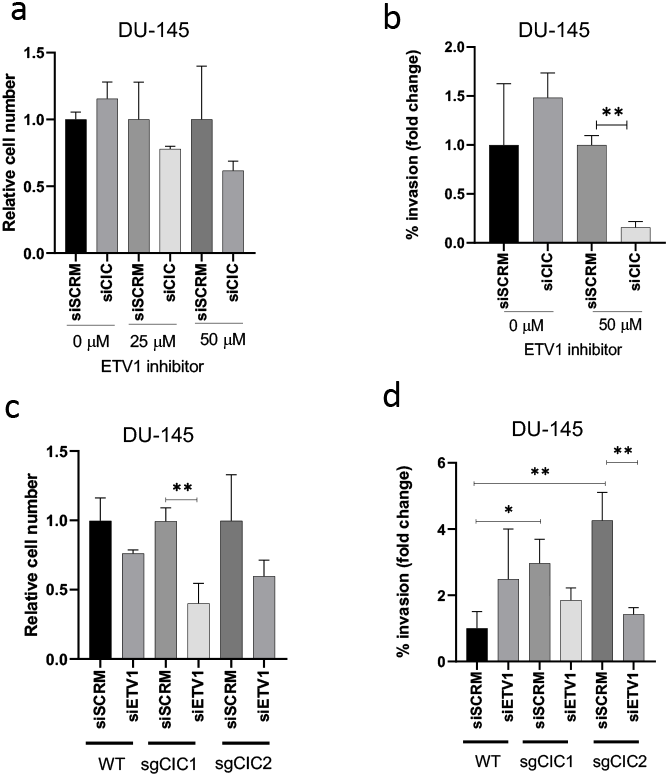
Combinatorial CIC and ERF loss leads to increased dependence on ETV1. (a, b) DU-145 cells were transfected with either siScramble or siCIC. After 48 hours, different concentrations (0 μM, 25 μM or 50 μM) of BMS32048 (ETV1 inhibitor) was added to both the transfected groups. Crystal violet viability (a) and transwell invasion assays (b) performed 24 hours after the addition of BMS32048. P value = **p<0.01. All siCIC groups were normalized to their respective siScrambled groups (taken as 1). (c, d) DU-145 WT and DU-145 CIC KO cells were transfected with either siScramble or siETV1 for 48 hours. Crystal violet viability (b) and invasion assays (d) were used to compare the different groups. P value = *p<0.05, **p<0.01. Error bars represent SD.

## Discussion

Molecular and functional subclassification of human prostate cancer has revealed a dependence on ETS family transcription factors including ERG, ETV1, ETV4, and ETV5 ^36,40^. The predominant mode of ETS activation in prostate cancer is through chromosomal rearrangements that fuse *ERG, ETV1, ETV4, and ETV5* to the androgen-regulated *TMPRSS2* gene, leading to fusion oncoproteins that drive oncogenesis ^40–44^. Interestingly, recent data indicate that ETV1, ETV4, and ETV5 are upregulated in a fusion-independent manner and are associated with poor clinical outcomes in prostate cancer patients ^3,45^. Our study focused on understanding the molecular mechanisms that drive fusion-independent upregulation of ETS family members and we reveal a new molecular subclass of prostate cancer defined by a co-deletion of two transcription factors, CIC and ERF.

CIC is a transcription factor that directly silences *ETV1, ETV4, and ETV5* transcription through direct repression at proximal regulatory sites ^21^. We observed that in ∼10-12% of human prostate cancer, CIC and ERF are co-deleted through focal homozygous or heterozygous deletions. It has been recently shown that ERF competes for ETS DNA binding motifs and our studies identify cooperative regulation of key target genes between CIC and ERF. Specifically, through ChIP-Seq analysis coupled with a series of *in vitro* and *in vivo* studies, we identify a coordinated binding of CIC and ERF to the proximal *ETV1* regulatory element that physically and functionally regulates ETV1 expression. Therefore, we reveal *ETV1* as a novel ERF target gene. Rescuing ERF in CIC deficient prostate cancer cells decreases *ETV1* expression and limits malignant phenotypes including viability, migration and invasion.

The 19q13.2 locus contains CIC and ERF, which are physically adjacent and oriented in opposing directions. The long and short isoforms of *CIC* are separated from *ERF* by ∼15 and 30kb, respectively. Thus, future studies directed at defining the genome topology and epigenetic states, both within and around this highly conserved region are warranted. In particular, studies aimed at mapping topology associated domains and key histone marks can potentially reveal shared upstream regulatory elements including enhancers or super-enhancers that may co-regulate CIC and ERF in concert. These findings could reveal non-genetic mechanisms to functionally regulate CIC and ERF expression in a coordinated fashion.

The use of ETV1 inhibitors has been limited to preclinical studies ^39^. These studies have largely focused on direct targeting of ETV1 fusion oncoproteins in prostate cancer ^39^. Our findings indicate that patients with CIC-ERF deficient prostate cancer may have a survival dependence on ETV1 and that these patients may indeed benefit from ETV1 directed therapies. Thus, the future development and clinical application of ETV1 inhibitors in patients that harbor the CIC-ERF co-deletion may be a viable therapeutic strategy. Furthermore, targeting the unique transcriptional program regulated by either CIC or ERF is another research direction for CIC-ERF co-deletion sub-population. Collectively, we have uncovered a molecular subset of prostate cancer defined by a co-deletion of CIC and ERF and further demonstrate a mechanism-based strategy to limit tumor progression through ETV1 inhibition in this subset of human prostate cancer.

## Materials and Methods

### Cell Lines, Drug, and Reagents

Cell lines were cultured as recommended by the American Type Culture Collection (ATCC). PNT2, DU-145 and PC-3 cells were purchased from ATCC. PNT2 ERF KD (shERFA1, shERFB1), PNT2 CIC KO (sgCIC1, sgCIC2) and PNT2 ERF KD+CIC KO were derived from parental PNT2 cells. shRNAs targeting ERF to develop PNT2 shERFA1 and PNT2 shERFB1 were obtained from Sigma-Aldrich: TRCN000001391, TRCN0000013912. Puromycin (1μg/ml) was used as a selection reagent. Two sgRNAs targeting CIC were previously validated and were gifts from William Hahn, Addgene (#74959 and #74953). These sgRNAs were used to develop PNT2 sgCIC1 and PNT2 sgCIC2 cells. Blasticidin (10μg/ml) was used as a selection agent. PNT2shERFA1+sgCIC1 and PNT2shERFA1+sgCIC2 were developed from the combination of the above two shRNA and sgRNAs. All PNT2 cells were grown in DMEM media supplemented with 10% FBS, 100 IU/ml penicillin and 100 μ g/ml streptomycin.

DU-145 ERF, DU-145 CIC KO (sgCIC1, sgCIC2) and DU-145 ERF +CIC KO (ERF+sgCIC1, ERF+sgCIC2) were derived from parental DU-145 cell line. Lentiviral GFP-tagged ERF (GeneCopoeia, EX-S0501-Lv122) was used to develop DU-145 ERF cells with puromycin (1μg/ml) as a selection marker. The above-mentioned two sgRNAs were used to develop DU-145 sgCIC1 and DU-145 sgCIC2 cells with blasticidin (15μg/ml) as the selection agent. DU-145 ERF +sgCIC1 and DU-145 ERF +sgCIC2 were developed from the combination of the above two.

PC-3 ERF KD (shERFA1), PC-3 ERF OE, PC-3 CIC OE, PC-3 (ERF+CIC) OE cells were derived from parental PC-3 cell line. shRNAs targeting ERF (Sigma-Aldrich: TRCN000001391) and lentiviral GFP-tagged ERF (GeneCopoeia, EX-S0501-Lv122) were used to develop PC-3 shERFA1 and PC-3 ERF OE respectively. PC-3 CIC OE cells were developed using CIC-Myc-tag plasmid purchased from Origene (CAT#: RC215209). Geneticin (250μg/ml) was used as a selection agent. PC-3 (ERF+CIC) OE cells were developed using a combination of ERF-GFP and CIC-Myc overexpressing plasmid.

All DU-145 and PC-3 cells were grown in RPMI 1640 media supplemented with 10% FBS, 100 I U/ml penicillin and 100 μ g/ml streptomycin, respectively. All cell lines were maintained at 37 °C in a humidified atmosphere at 5% CO2.

BRD32048 is an ETV1 inhibitor that was purchased from Sigma-Aldrich (CAT#: SML1346).

### Subcutaneous Tumor Xenograft Assays

Four week old male SCID mice were purchased from Jackson Laboratory. Mice were kept under specific pathogen-free conditions and facilities were approved by the UCSF IACUC. To prepare cell suspensions, PNT2 and its other genetic variants (PNT2 ERF KD, PNT2 CIC KO, PNT2 ERF KD+CIC KO) were briefly trypsinized, quenched with 10% FBS RPMI media and resuspended in PBS. Cells were pelleted again and mixed with PBS/Matrigel matrix (1:1) for a final concentration of 0.1×10^5^ cells/µl. A 100 μl cell suspension containing 1 × 10^6^ cells were injected (s.c.) in the right and left flanks of immunodeficient mice (n=10/group). Mice were observed for tumor formation in different groups over a few weeks.

For other subcutaneous xenografts, four week old male mice (NU/J) were purchased from Jackson Laboratory and were six-eight weeks old at time of experiment. 1.0 × 10^6^ DU-145 cells and its variants (DU-145 ERF, DU-145 CICKO and DU-145 ERF +CICKO) were resuspended in PBS/Matrigel (1:1) matrix and injected s.c. into the right and left flanks of nude mice (n=10/group). Tumor volume was measured twice per week using Vernier caliper. Tumor volume was determined using caliper measurements of tumor length (L) and width (W) according to the formula V = (L X W2) X 0.52.

Similarly, 1.0 × 10^6^ PC-3 tumor cells and its variants (PC-3 ERFOE, CICOE) were injected subcutaneously in flanks of male nude mice (NU/J), n=10/group and tumor volume was monitored in different groups. Mice body weight was measured in all the experiments throughout the study. At the end of the experiment, mice were sacrificed by CO2 overdose in accordance with IACUC guidelines.

### Gene Knockdown and Overexpression Assays

ON-TARGET plus Scramble, ETV1 (L-003801-00-0005) and CIC (L-015185-01-0005) siRNAs were obtained from GE Dharmacon and transfection was performed with Dharmafect transfection reagent per manufacturer recommendations. ETV1 inhibitor-BRD32048 (SML1346) was purchased from Millipore Sigma. Lentiviral GFP-tagged ERF was obtained from GeneCopoeia (EX-S0501-Lv122). pCMV-CIC with myc-tag was purchased from Origene and validated previously.

### Real-Time quantitative PCR

Isolation and purification of RNA was performed using RNeasy Mini Kit (Qiagen). 500 ng of total RNA was used in a reverse transcriptase reaction with the SuperScript III first-strand synthesis system (Invitrogen). Quantitative PCR included three replicates per cDNA sample. Human CIC, ERF, ETV1, ETV4, ETV5, and endogenous controls GAPDH were amplified with Taqman gene expression assay (Applied Biosystems). Expression data were acquired using an ABI Prism 7900HT Sequence Detection System (Thermo Fisher Scientific). Expression of each target was calculated using the 2−ΔΔCt method and expressed as relative mRNA expression.

### Western Blot Analysis

Adherent cells were washed and lysed with RIPA buffer supplemented with proteinase and phosphatase inhibitors. Proteins were separated by SDS-PAGE, transferred to Nitrocellulose membranes, and blotted with antibodies recognizing: CIC (Thermo Fisher Scientific –PA146018), ERF (Thermo Fisher Scientific –PA530237), ETV1 (Thermo Fisher Scientific –MA515461), HSP90 (Cell Signaling– 4874S), Actin (Cell Signaling – 4970S). All immunoblots represent at least two independent experiments.

### Luciferase Promoter Assay

293T and DU-145 cells were split into a 96 well plate to achieve 50-70% confluence the day of transfection. LightSwitch luciferase assay system (SwitchGear Genomics) was used per the manufacturer’s protocol. Briefly, a mixture containing FuGENE 6 transfection reagent, 50ng Luciferase GoClone ETV1 promoter (S720645) plasmid DNA, 50ng of either control (empty) vector or fully sequenced ERF cDNA (GeneCopoeia (EX-S0501-Lv122)), were added to each well. All transfections were performed in quintuplicate. The plates were assessed for luciferase activity after 48 hours of treatment.

### Chromatin Immunoprecipitation with Sequencing (Chip-Seq) And PCR

ChIP was performed on PNT2 and DU-145 cells with the SimpleChIP Enzymatic Chromatin IP kit, Cell Signaling Technology #9003 in accordance with the manufacturer’s protocol. The antibodies used for IP were as follows: CIC (Thermo Fisher Scientific – PA146018) and ERF (Thermo Fisher Scientific –PA530237). Paired-end 150-bp (PE150) sequencing on an Illumina HiSeq platform was then performed. ChIP-Seq peak calls were identified through Mode-based Analysis of ChIP-Seq (MACS). For ETV1 ChIP-PCR validation, primers were designed in the proximal regulatory element of ETV1. The promoter primer sequences are as follows:

ETV1-CIC-Forward-1-(5’ CAGGACAAAGAGGAGGCAGCGAGCTG-3’)

ETV1-CIC-Reverse-1-(5’ GTTTATTGCTGACCCCTCAGCGCTCCGC 3’)

ETV1-ERF-Forward-1-(5’-CCAGGTCCGGGGTTGAGTGCTGTGC-3)

ETV1-ERF-Reverse-1 (5’-CATTTGTGACCAGAACTAGTGACC-3)

VCaP ERF ChIP-Seq was previously performed (Bose et al., Nature 2017) and publicly available in the GEO database: GSE83653. For this analysis we used the following samples:

GSM2612450_VCaP_red_r1_peaks

GSM2612451_VCaP_red_r2_peaks

GSM2612454_INPUT red replicate one

GSM2612455_INPUT red replicate two

### Colony formation Assay

Equal number of cells from different groups (500–600 cells/well) were seeded in a 6-well plate. Cells were allowed to form colonies for 7 days. At day 7, cells were fixed and stained with 0.5% crystal violet solution after washing with PBS and performed in triplicate. Finally, the colonies with >50 cells were counted under an image J-software.

### Tumorsphere Assay

Approximately 25,000 cells from different groups were cultured in tumorsphere media at 37°C and 5% CO2 for 7 days. Tumorsphere medium contains serum free DMEM /F12 supplemented with 10 ng/ml FGF (fibroblast growth factor), 20 ng/ml EGF (epidermal growth factor), 1xITS (Insulin-Transferrin-Selenium) and B27 supplement. On day 7, images of different areas of the wells were taken using confocal. The size of the sphere was calculated using Fiji (Image J) software in all the tested groups. Each group consisted of three replicate wells and at least 6 images (n>/=6).

### Viability Assays

5000 cells were plated in a 12 well plate. Crystal violet staining was performed 5 days after cell plating with 3 replicates per group. CellTiter Glo experiments were performed according to the manufacturer’s protocol. In brief, cells were plated in a 96-well plate, and analyzed on a Spectramax microplate reader (Molecular Devices) at different days. Each assay was performed with 6 replicate wells.

### Transwell invasion assays

RPMI with 10% FBS was added to the bottom well of a trans-well chamber. 2.5×10^4 cells resuspended in serum free media was then added to the top 8 µm pore matrigel coated (invasion) or non-coated (migration) trans-well insert (BD Biosciences). After 20 hours, non-invading cells on the apical side of inserts were scraped off and the trans-well membrane was fixed in methanol for 15 minutes and stained with Crystal Violet for 30 minutes. The basolateral surface of the membrane was visualized with a Zeiss Axioplan II immunofluorescent microscope at 5×. Each trans-well insert was imaged in five distinct regions at 5× and performed in triplicate. % invasion was calculated by dividing the mean # of cells invading through Matrigel membrane / mean # of cells migrating through control insert.

### Wound healing assays

Cells were plated at a density of 0.5×10^6^ cells/well and incubated to form a monolayer in 6-well dishes. Once a uniform monolayer was formed, wound was created by scratching the monolayer with a 1ml sterile tip. Floating cells were removed by washing the cells with PBS three times. Images of the wound were taken at this point using bright field microscope and considered as a 0 hour time point. Further, media was added in all the wells and cells were left to migrate either for 24 hours (DU-145 cells) or 48 hours (PNT2 ond PC-3 cells). At end point, wound was imaged again using bright field microscope. The wound area at different points was calculated using Image J software. Each group consisted of at least three replicate wells.

### Analysis of prostate cancer datasets from cBioPortal

15 prostate cancer datasets (see figure 1c for individual studies) were queried for alterations in CIC and ERF using the cBioPortal platform “query by gene” function. Stratification into “ERF-CIC No co-deletion” and “ERF-CIC co-deletion” was performed in cBioPortal and associated with “Gleason Score”, “AJCC Primary Tumor T Stage”, and Disease-free and Progression-free survival using the “Plots” and “Comparison/Survival” functions. P-values for comparison between Gleason and Tumor Stage were calculated using Fisher’s exact test and survival curves were calculated by Log-rank test.

For the CIC and ERF mutational analysis in primary prostate versus metastatic castrate resistant prostate cancer, we selected studies that purely represented primary prostate tumors (PNAS 2014, Cell 2014, Nature 2017) and advanced mCRPC (Nature 2012, Cell 2015, PNAS 2019).

### DAVID Functional Clustering Analysis

The 91 shared candidate target genes between CIC and ERF identified through our ChIP-Seq analysis in PNT2 and VCaP cells, respectively, were used as an input list for analysis using the DAVID Bioinformatics Database.

### Single-sample Gene Set Enrichment Analysis (ssGSEA)

Gene-level expression of samples are computed using RSEM (PMID: 18505969) (Version 1.3.3) and log-2 normalized. ssGSEA analysis is conducted using the ssGSEA module (Version 10.0.9) on GenePattern (PMID: 16642009) for selected established signature gene sets. ssGSEA result is then processed and information coefficient and statistical significance levels are computed among signature gene sets.

### Statistical analysis

Experimental data are presented as mean +/-Standard Deviation (SD) or Standard Error of the Mean (SEM). P-values derived for all *in-vitro* experiments were calculated using two-tailed student’s t-test or one-way ANOVA. The detailed statistical analysis performed for each experiment is defined in the figure legends.

### Study approval

For tumor xenograft studies, specific pathogen-free conditions and facilities were approved by the American Association for Accreditation of Laboratory Animal Care. Surgical procedures were reviewed and approved by the UCSF Institutional Animal Care and Use Committee (IACUC), protocol #AN178670-03.

## Acknowledgements

The authors thank members of the Okimoto and Huang labs and funding support from the following sources: A National Cancer Institute (NCI) Cancer Center Support Grant (P30CA082103) and Benioff Initiative for Prostate Cancer Research Awards (N.G and R.A.O), a NIGMS Predoctoral Training in Biomedical Sciences T32 (C.L., T32 GM 136547), Basic Science Research Program through the National Research Foundation of Korea (NRF) funded by the Ministry of Education (NRF-2020R1A6A3A03039483)(J.W.K) and NCI K08-CA222625 and R37CA255453 awards to R.A.O.

## Author information

Department of Medicine, University of California, San Francisco, San Francisco California, USA

Nehal Gupta, Hanbing Song, Wei Wu, Rovingaile Kriska Ponce, Yone Kawe Lin, Ji Won Kim, Eric J Small, Franklin W. Huang, and Ross A. Okimoto

Department of Radiation Oncology, University of California, San Francisco, San Francisco California, USA

Felix Y. Feng

Helen Diller Family Comprehensive Cancer Center, University of California, San Francisco, San Francisco, California, USA

Eric J Small, Felix Y. Feng, Franklin W. Huang, and Ross A. Okimoto

## Ethics declarations

The authors have no competing interest and have nothing to declare.

---

**Supplementary Figure 1.**
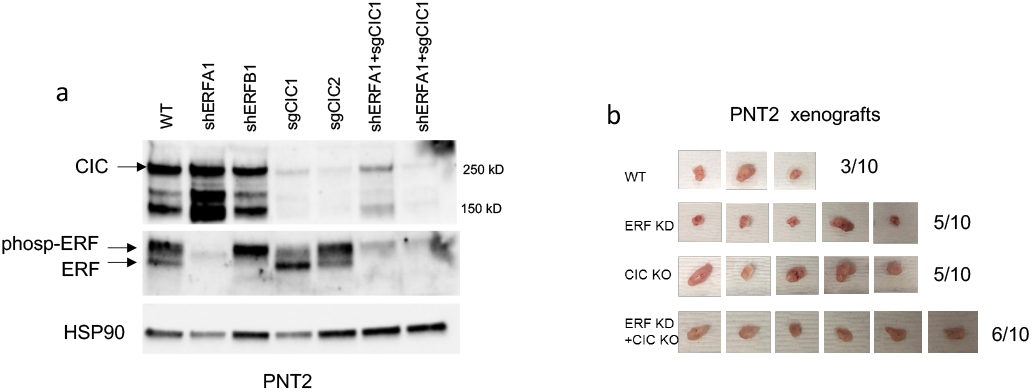
CIC and ERF loss enhances tumor formation in PNT2 cells. (a) Immunoblot of CIC, ERF and HSP90 in PNT2 and its different variants. Representative figure; performed in duplicate. Arrows indicate CIC and ERF bands. (b) Tumor explants from mice in figure 2E.

**Supplementary Figure 2.**
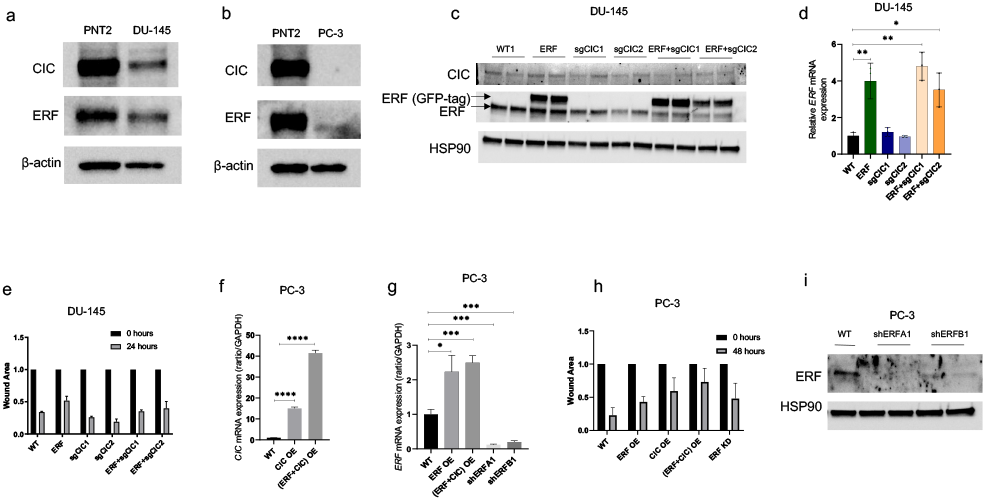
DU-145 and PC-3 prostate cancer cells are well defined model systems to study CIC and ERF function. (a) Immunoblot of CIC, ERF, and beta-actin in DU-145 cells compared to PNT2 cells. (b) Immunoblot of CIC, ERF, and beta-actin in PC-3 cells compared to PNT2 cells. (c) Immunoblot of CIC, ERF and HSP90 in parental DU-145 cells and engineered DU-145 different variants. Representative figure; performed in duplicate. (d) Relative ERF mRNA expression in DU-145 parental, ERF OE, CIC KO, and ERF OE+CIC KO, Student’s t-test, ***p<0.001, ****p<0.0001. (e) Wound healing assay in DU-145 cells with ERF rescue, CIC KO, or ERF rescue + CIC KO compared to parental. (f) CIC mRNA expression in PC-3 parental, PC-3 CIC OE, and PC-3 (ERF+CIC) OE. P value = ****p<0.0001 (g) Relative ERF mRNA expression in PC-3 parental, PC-3 ERF OE, PC-3 (ERF+CIC) OE, and PC-3 ERF KD. P value = *p<0.05, **p<0.01, ***p<0.001. (h) Wound healing assay in PC-3 cells expressing ERF OE, CIC OE, (ERF+CIC) OE, or ERF KD compared to control. (i) immunoblot of ERF and HSP90 in PC-3 cells with ERF KD.

**Supplementary Figure 3.**
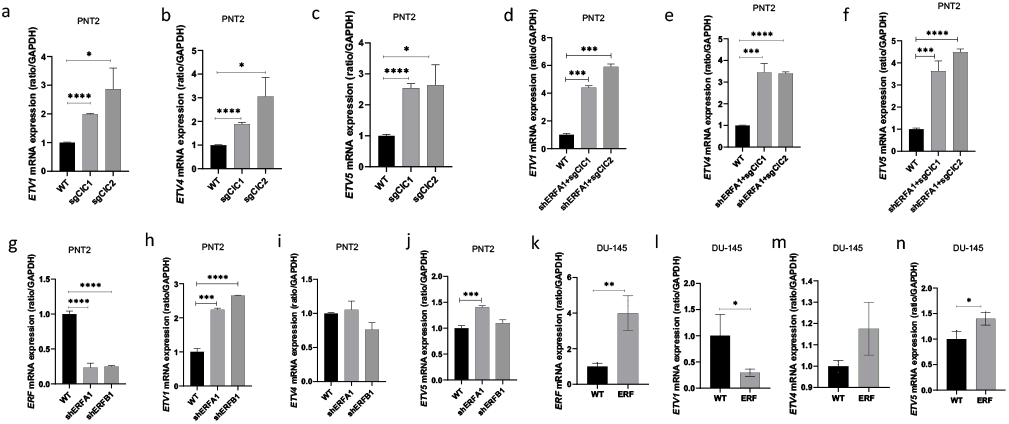
ETV1, but not ETV4 or ETV5, is a transcriptional target of both ERF and CIC. (a-c) Relative ETV1, ETV4 and ETV5 mRNA expression in PNT2 parental and PNT2 CIC KO cells. (d-f) Relative ETV1, ETV4 and ETV5 mRNA expression in PNT2 parental and PNT2 ERF KD+CIC KO cells. (g) Relative ERF, (h) ETV1, (i) ETV4, and (j) ETV5 mRNA expression in parental PNT2 and PNT2 ERF KD cells. Relative (k) ERF, (l) ETV1, (m) ETV4, and (n) ETV5 mRNA expression in parental DU-145 and DU-145 ERF cells. P values for all figures = *p<0.05, **p<0.01, ***p<0.001, ****p<0.0001.

**Supplementary Figure 4.**
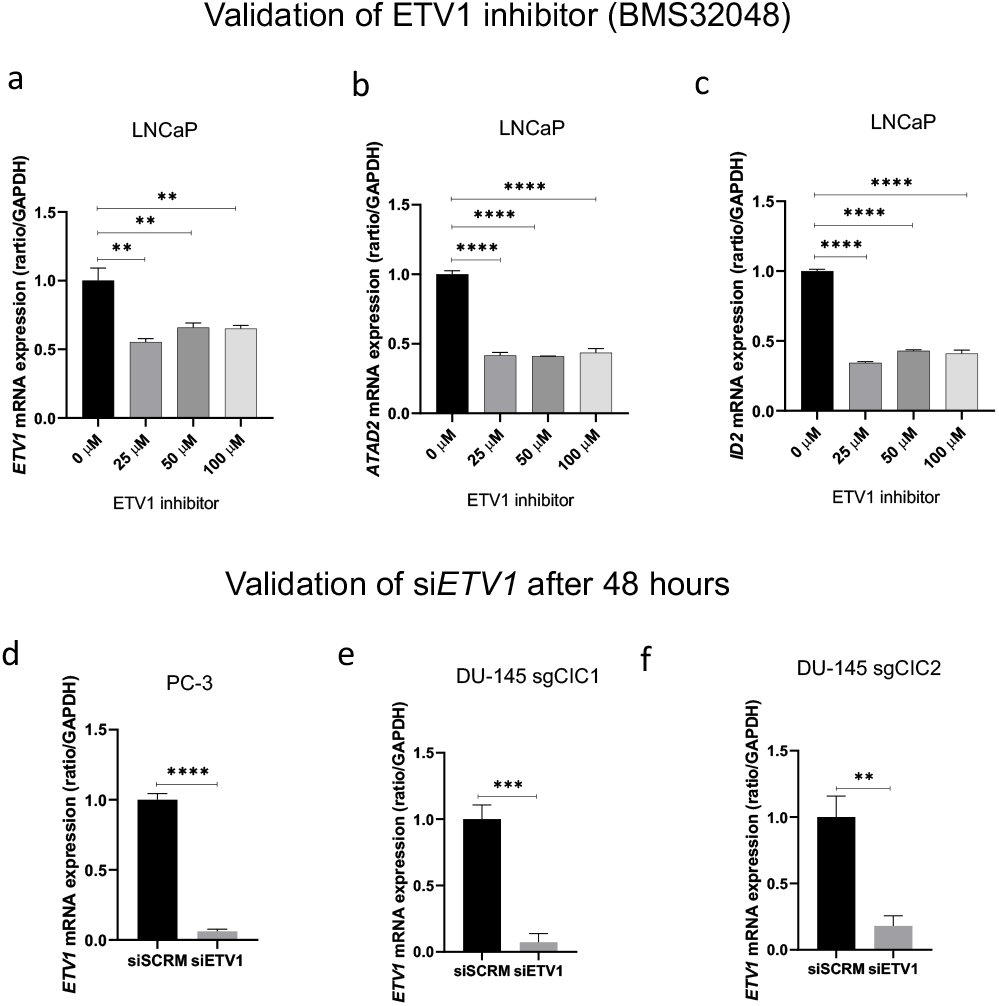
Validation of ETV1 chemical (BMS32048) and genetic inhibition in prostate cancer cells. (a-c) Relative ETV1, ATAD2 and ID2 mRNA expression in LNCaP cells with or without the BMS32048 (ETV1 inhibitor) treatment. (d) Relative ETV1 mRNA expression in PC-3 cells, (e) DU-145+sgCIC1 and (f) DU-145+sgCIC2 with or without siETV1. P values for all figures = **p<0.01, ***p<0.001, ****p<0.0001.

